# Lysosomal Profiling with LysoTracker for Quantitative Assessment of Cellular Senescence in Human Fibroblasts

**DOI:** 10.64898/2026.04.16.719007

**Authors:** Javier Estrada, Scott Tenenbaum, Melinda Larsen, Thomas J Begley, J.A. Melendez

## Abstract

Cellular senescence is a stable cell-cycle arrest state associated with characteristic phenotypes, including enlarged cell morphology, altered secretory signaling, and pronounced lysosomal remodeling. Senescent cells commonly accumulate increased numbers of enlarged lysosomes with changes in acidity and degradative capacity, creating an opportunity for simple live-cell readouts of senescence-linked organelle remodeling. Here, I describe a live-cell lysosomal profiling protocol that uses LysoTracker Deep Red, an acidotropic fluorescent dye, to label and quantify acidic organelles in individual living cells as an indicator of senescence-associated lysosomal expansion. The method is demonstrated in IMR-90 human lung fibroblasts undergoing replicative senescence across serial passaging. The protocol details cell culture and passage tracking, LysoTracker staining, fluorescence imaging, and straightforward image-based quantification of lysosomal signal intensity and lysosome-enriched area per cell. As an optional validation step, senescence-associated β-galactosidase staining is performed on parallel cultures to confirm senescent cell identity. Representative outcomes show increased LysoTracker signal and expanded lysosome-enriched regions in late-passage cultures compared to early-passage controls, consistent with lysosomal remodeling during senescence. This protocol is designed to be simple to adopt and can be adapted to other cell types or senescence-inducing stresses, providing a practical, quantitative complement to conventional endpoint assays.

**SUMMARY:** This article presents a live-cell imaging protocol using LysoTracker Deep Red to quantify lysosomal remodeling as a marker of cellular senescence in IMR-90 human fibroblasts. We demonstrate quantitative lysosomal readouts derived from fluorescence imaging, including lysosome-enriched area and intensity measurements that can be summarized per cell and, when desired, as stitched-field, per-nucleus normalized metrics. Senescence status can be validated against senescence-associated β-galactosidase (SA-β-Gal) staining performed on parallel cultures. The method can be adapted to other cell types or senescence-inducing stresses and enables quantitative analysis of lysosomal remodeling during senescence.

## INTRODUCTION

Cellular senescence is a durable cell-cycle arrest program activated by various stresses and aging-related triggers.^1–4^ Senescent cells are classically identified by a constellation of markers, since no single marker is completely specific. Hallmarks include enlarged, flattened cell morphology, senescence-associated β-galactosidase (SA-β-Gal) activity, formation of DNA damage foci, and a pro-inflammatory secretory profile (SASP).^2,4,5^ Among these features, lysosomal remodeling has emerged as a prominent and functionally significant trait of senescence.^6,7^ Senescent cells display *profound lysosomal changes*, notably a dramatic expansion in lysosomal size and number along with altered function (often including partial neutralization of lysosomal pH and accumulated undegraded material).^6–9^ This lysosomal expansion contributes to the increase in SA-β-Gal activity, an established senescence marker that reflects elevated lysosomal β-galactosidase content in senescent cells.^10–12^ Indeed, SA-β-Gal staining exploits this phenomenon by using a suboptimal pH (6.0) incubation to selectively precipitate dye in senescent-cell lysosomes, which have accumulated the enzyme and exhibit a higher pH than young cells.^10,12,13^

Lysosomes play central roles in protein turnover and signaling, and their dysregulation is now considered one of the hallmarks of aging and senescence.^1,2,4,6^ Senescent cells often have **dysfunctional lysosomes**, evidenced by elevated luminal pH, altered enzyme activity, and accumulation of lipofuscin (an autofluorescent pigment of oxidized macromolecules).^6,14^ The biogenesis of lysosomes is frequently upregulated or altered via the coordinated lysosomal expression and regulation network (CLEAR) under control of transcription factor EB (TFEB) and related factors.^6,7,15^ This can lead to an **increased lysosomal mass** in senescent cells, as the cell adapts to stress by expanding its degradative compartment.^6,7,10^ Conversely, senescent cells may experience lysosomal functional decline (e.g., less acidic pH), which can impact processes like autophagy and render cells resistant to certain stresses such as ferroptotic cell death.^6,8^ Thus, measuring lysosomal content and function is key to understanding and identifying senescence.^6,16^

**LysoTracker Deep Red** is a membrane-permeable fluorescent dye that selectively accumulates in acidic organelles (late endosomes and lysosomes) due to protonation in low pH environments.^16^ Its far-red emission spectrum makes it suitable for live-cell imaging with minimal overlap with cellular autofluorescence.^16^ The intensity of LysoTracker staining in a cell correlates with the volume and surface area of its lysosomes, providing an indirect measure of organelle abundance and size. Prior studies have shown that senescent cells exhibit stronger LysoTracker fluorescence than non-senescent cells, consistent with their higher lysosomal content.^9^ By quantifying LysoTracker fluorescence at the single-cell level, one can assess lysosomal biogenesis and morphology changes associated with senescence.^6,16^ This live-cell approach has several advantages over the conventional SA-β-Gal assay: it enables *quantitative* and *spatially resolved* analysis of lysosomes in individual living cells, can be combined with other fluorescent reporters, and does not require cell fixation (allowing subsequent functional assays on the same cells if needed). Moreover, it captures heterogeneity in senescent cell populations by measuring continuous features (lysosomal size, count, intensity) rather than a binary staining endpoint.

In this protocol, we detail the use of LysoTracker Deep Red to profile lysosomal changes in IMR-90 fibroblasts undergoing replicative senescence. IMR-90 is a normal human diploid fibroblast line (female, fetal lung origin) that is widely used in aging research; like other primary-like fibroblasts (e.g., WI-38), IMR-90 cells enter senescence after ∼50–60 population doublings (the Hayflick limit) due to telomere shortening and DNA damage signaling. We describe how to culture IMR-90 cells to induce senescence via serial passaging, verify senescence by morphology and SA-β-Gal staining, and then stain live cultures with LysoTracker Deep Red for fluorescent imaging.^5,10,12,13^ The protocol covers image acquisition and analysis, including how to measure lysosomal count per cell, total lysosomal area per cell, and mean fluorescence intensity as quantitative readouts. We also emphasize experimental controls and safety: for example, careful handling of dyes and fixatives, minimizing phototoxicity during imaging, and accounting for cellular autofluorescence in aged cells. The method is demonstrated on replicative senescence, but it can be adapted to other models such as drug-induced senescence (e.g., doxorubicin or etoposide treatment) or stress-induced premature senescence (SIPS) by oxidants.^2–4^ In each case, the LysoTracker-based readouts should be compared to appropriate controls (proliferating or quiescent cells) and, if possible, cross-validated with at least one traditional senescence marker (such as SA-β-Gal or p16INK4a expression).^3,5,12,13^

This live-cell lysosomal profiling fills a practical gap in senescence assays: it provides a reproducible, single-cell resolution measurement of a key organelle phenotype of senescence, lysosomal expansion, which is directly tied to senescent cell function (e.g., enhanced degradative capacity and secretory activity).^2–4,6^ Because LysoTracker staining is compatible with high-throughput imaging and flow cytometry, the protocol can be scaled up for drug screening (e.g., testing senolytics or modulators of lysosomal function).^2,3,6^ In the *Discussion*, we additionally outline how one could integrate an AI-based classifier trained on lysosomal features to automatically identify senescent cells in images, as an optional extension for laboratories interested in computational approaches. Overall, this method allows investigators to visualize and quantify the lysosomal dimension of senescence biology, shedding light on how lysosomal biogenesis and morphology are altered as cells age or undergo stress, and providing a platform for testing interventions that target the lysosome in senescent cells. The image analysis workflow used to segment nuclei, derive cell masks, quantify LysoTracker features, and optionally generate SA-β-Gal-based labels is implemented in an open-source companion package (SenTrack) provided as Supplemental Files and maintained at: https://github.com/goldbader-hub/Sentrack

## PROTOCOL

Experiments use an established human cell line (IMR-90). Perform all work in compliance with institutional biosafety guidelines and local regulations for handling human-derived cell lines.

### 1. General preparation and workflow overview

1.1 Define the experimental goal as a live-cell lysosomal profiling assay that uses LysoTracker Deep Red staining and a live nuclear dye to quantify lysosomal remodeling as a continuous readout of senescence burden in IMR-90 cells. Define the primary biological comparison for the experiment as one of the following designs:

1.1.1 Senescence induction vs matched non-induced control.

1.1.2 Senescence-high vs senescence-low conditions across a dose or time series.

1.1.3 Senescence-high condition treated with a senolytic vs vehicle-treated senescence-high control.

1.1.4 Senolytic treatment series that includes both a senescence-low baseline group and a senescence-high group.

1.2 Establish the minimum control structure for all experiments.

1.2.1 Include a senescence-low control group that matches plating density, staining timing, and imaging settings.

1.2.2 Include a senescence-high group created by a defined senescence trigger or by late-passage replicative senescence.

1.2.3 Include biological replicates per condition and perform staining and imaging using identical timing across conditions.

1.3 Define the assay readouts as direct image-based measures of lysosomal remodeling:

1.3.1 Measure per-cell mean LysoTracker intensity, integrated LysoTracker intensity, and lysosome-enriched area per cell using a consistent intensity threshold within the cell-associated region.

1.3.2 Record optional lysosome puncta count and size metrics when imaging resolution supports puncta discrimination.

1.4 Use senescence-associated β-galactosidase (SA-β-Gal) staining as an optional validation step performed on parallel wells or matched samples, depending on experimental constraints. Use this validation to confirm that late-passage cultures show increased SA-β-Gal positivity relative to early-passage cultures.

1.5 Standardize the assay across conditions by fixing the following experimental variables prior to data collection:

1.5.1 LysoTracker working concentration (75 nM) and staining duration (30 min).

1.5.2 Nuclear dye identity and working concentration range (for example, Hoechst 33342 at 1 µg/mL to 2 µg/mL).

1.5.3 Imaging parameters including objective magnification, exposure time, illumination intensity, binning, and focus strategy.

1.5.4 Plate format and imaging configuration (clear-bottom imaging plates or glass-bottom dishes), including replicate well layout.

1.6 Acquire images under conditions that preserve live-cell signal quality and reduce technical variability. Keep early- and late-passage plates protected from light during staining and transport and keep the time outside controlled culture conditions as short as possible during imaging sessions.

1.7 Save fluorescence images as single-plane, single-channel, lossless TIFF files for each channel. Use a consistent naming scheme that preserves pairing between nuclear and LysoTracker images and keeps fields traceable to plate, well, condition, and replicate. Use a shared base name for the paired channels and add channel suffixes (for example, _DAPI.tif or _Hoechst.tif for the nuclear image and _Lyso.tif for the LysoTracker image). Store all images for a single experiment in a dedicated folder with subfolders organized by plate or imaging date if needed.

NOTE: A practical pause point is after establishing early- and late-passage cohorts and confirming that late-passage cultures show slowed growth and senescence-like morphology.

1.8 Optional: Include oxygen as an experimental variable by maintaining matched IMR-90 cohorts in both ambient oxygen and low oxygen. Use matched early- and late-passage groups under each oxygen condition and process all groups identically through staining and imaging.

### 2. Cell culture and induction of replicative senescence

2.1 Thaw IMR-90 human lung fibroblasts according to standard cell culture practices and expand cells in Minimum Essential Medium Eagle supplemented with 10% fetal bovine serum. Add penicillin-streptomycin at 1% if consistent with laboratory practice and experimental design. Maintain cells at 37 °C in 5% CO_2_.

2.2 Maintain cultures in logarithmic growth and passage before 70% to 80% confluence. Use consistent media handling and feeding schedules to reduce variability across experimental runs.

2.3 Define the senescence-low baseline group as actively proliferating IMR-90 cultures maintained under conditions that preserve normal growth behavior. Match plating density and handling to all experimental groups.

2.4 Generate a senescence-high group using one of the following approaches, chosen based on experimental aims.

2.4.1 Replicative senescence by serial passaging with population doubling tracking until slowed proliferation and senescence-associated morphology are evident.

2.4.2 Stress-induced senescence by applying a defined senescence trigger and allowing an appropriate phenotypic development window.

2.4.3 Treatment-induced senescence by exposing cells to the condition under study when the goal is to test whether that condition induces senescence.

2.5 Validate that the selected approach produces a senescence-high phenotype using at least one orthogonal readout performed on parallel wells. Perform SA-β-Gal staining as an optional validation method and document the fraction of positive cells per condition when included.

PAUSE POINT: Pause after establishing senescence-low and senescence-high cohorts and confirming a clear phenotypic difference by morphology and optional validation staining.

2.6 Optional: Senolytic treatment design

2.6.1 Plate senescence-low and senescence-high conditions in replicate wells and allow cells to adhere and recover prior to treatment.

2.6.2 Treat senescence-high wells with senolytic compound(s) and include a vehicle-treated senescence-high control.

2.6.3 Include a senescence-low group treated with the same vehicle and, when relevant, the same compound concentration(s) to monitor non-senescent sensitivity.

2.6.4 Proceed to live LysoTracker and nuclear staining at a consistent time after treatment across all conditions.

2.7 Monitor cultures during late-passage/treatment progression and document morphology and growth patterns. Identify senescent cultures by slowed proliferation, enlarged and flattened morphology, and increased cytoplasmic granularity. Once growth arrest is evident, maintain the culture as the senescence-high cohort for staining and imaging rather than passaging further.

2.8 Minimize confounding stressors during culture maintenance by using consistent media preparation, consistent feeding schedules, and consistent handling time outside the incubator. Treat temperature fluctuations, extended time outside controlled CO_2_, and inconsistent media warming as sources of avoidable technical variability.

2.9 Optional: Maintain matched cultures under ambient oxygen and low oxygen. Maintain cultures under 21% O_2_ and under 5% O_2_ as separate cohorts and keep each cohort under its assigned oxygen condition during routine culture handling when feasible. Limit exposure to ambient oxygen during transfers and imaging preparation for low oxygen cohorts.

NOTE: A practical pause point is after defining and stabilizing early-passage and late-passage cohorts and confirming that late-passage cultures show consistent slowed growth and senescence-associated morphology.

### 3. Live-cell lysosomal and nuclear staining with LysoTracker Deep Red and a nuclear dye

3.1 Plate IMR-90 cells for all experimental conditions in clear-bottom imaging plates or glass-bottom dishes. Include a senescence-low control condition and at least one senescence-high condition generated by the selected trigger, and include biological replicates for each condition.

3.2 Seed cells to achieve a subconfluent monolayer on the day of staining. Record the seeding density and the time between plating and staining for each experiment to support run-to-run comparability.

3.3 Warm complete culture medium to 37 °C. Prepare a LysoTracker Deep Red working solution at 75 nM in the warmed medium and protect it from light.

3.4 Prepare sufficient LysoTracker working solution for all wells that will be stained in the run. Mix the working solution by gentle inversion and keep it protected from light until use.

3.5 Aspirate spent medium from each well. Add LysoTracker working solution gently along the wall of the well to avoid disturbing the monolayer and maintain a consistent staining volume across wells.

3.6 Incubate cells with LysoTracker for 30 min at 37 °C in 5% CO_2_. Keep all experimental groups under the same incubation duration and temperature.

3.7 Add a live nuclear dye during the final 5 min to 10 min of LysoTracker incubation. Add Hoechst 33342 directly to the LysoTracker-containing medium at the manufacturer-recommended working concentration, for example 1 µg/mL to 2 µg/mL, and mix by gentle rocking of the plate.

3.8 Remove the staining medium at the end of the incubation. Rinse cells once with pre-warmed phosphate-buffered saline to reduce background fluorescence.

3.9 Add pre-warmed phenol red-free culture medium or an imaging buffer immediately after the rinse. Proceed directly to live imaging and keep the time between buffer exchange and image acquisition consistent across all wells in the experiment.

3.10 Protect plates from light during transfers and imaging setup. Minimize time outside controlled temperature and CO_2_ conditions during handling and image acquisition.

CAUTION: Handle fluorescent dyes using gloves and eye protection. Dispose of dye-containing liquids and contaminated consumables according to institutional chemical safety procedures.

NOTE: Use a pause point after Step 3.9 if imaging requires staging multiple plates. Keep plates protected from light at 37 °C and resume imaging as soon as possible to limit staining drift.

NOTE: If experimental design includes oxygen as a variable, maintain each cohort under its assigned oxygen condition during staining and incubation, and record the handling time outside the assigned condition.

### 4. SA-β-Gal staining for validation (optional)

4.1 Use SA-β-Gal staining as an optional validation step to confirm that the senescence-high condition shows increased SA-β-Gal activity relative to the senescence-low control. Perform validation on parallel wells processed in the same plate format and under the same handling timeline as the imaging wells.

4.2 Complete live imaging for the validation wells before fixation. When desired, the same field can be imaged live, then the well can be fixed and processed for SA-β-Gal to enable matched pre- and post-fixation comparisons (Figure 3).

**Figure 1.**
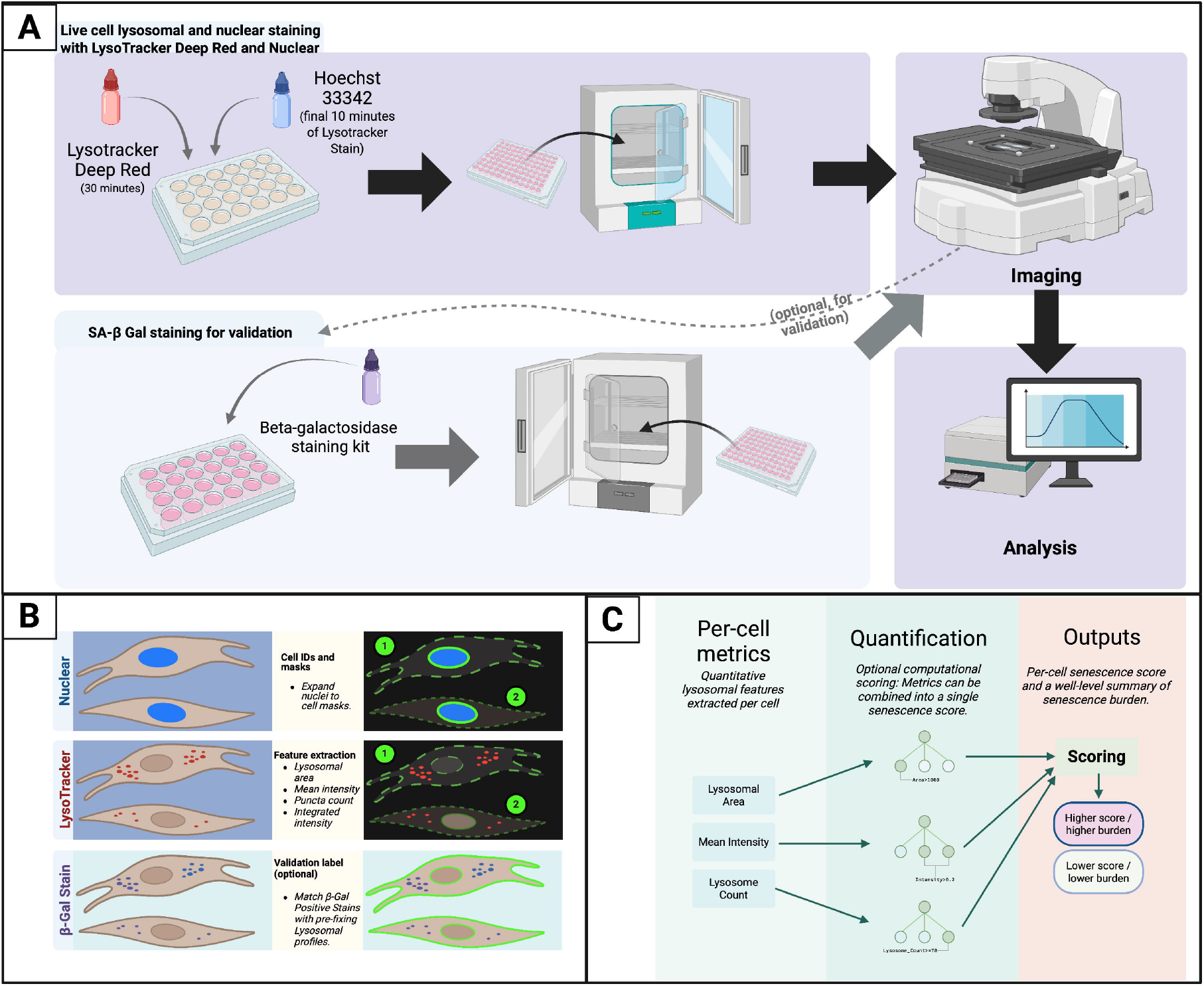
Schematic overview of a live-cell imaging workflow that quantifies senescence-associated lysosomal remodeling and summarizes senescence burden across conditions. **(A)** Experimental workflow. Cells are stained live with LysoTracker Deep Red and a nuclear dye, imaged by standard fluorescence microscopy, and optionally processed in parallel for SA-β-Gal staining for validation. **(B)** Image-derived per-cell measurements. The nuclear channel is used to identify individual cells, and nucleus-anchored expansion defines a per-cell region for extracting lysosome-associated measurements from the LysoTracker channel. Representative per-cell measurements include lysosomal area, mean LysoTracker intensity, and lysosome count. SA-β-Gal staining can be used as an optional validation label when generating or confirming reference datasets. **(C)** Quantification and outputs. Per-cell lysosomal measurements are combined into a single per-cell senescence score or probability using a simple scoring approach. Outputs include per-cell values and a well-level or condition-level summary of senescence burden that can be compared across senescence induction or senolytic treatment conditions. Created in BioRender. Estrada, J. (2026) https://BioRender.com/100ao08

**Figure 2.**
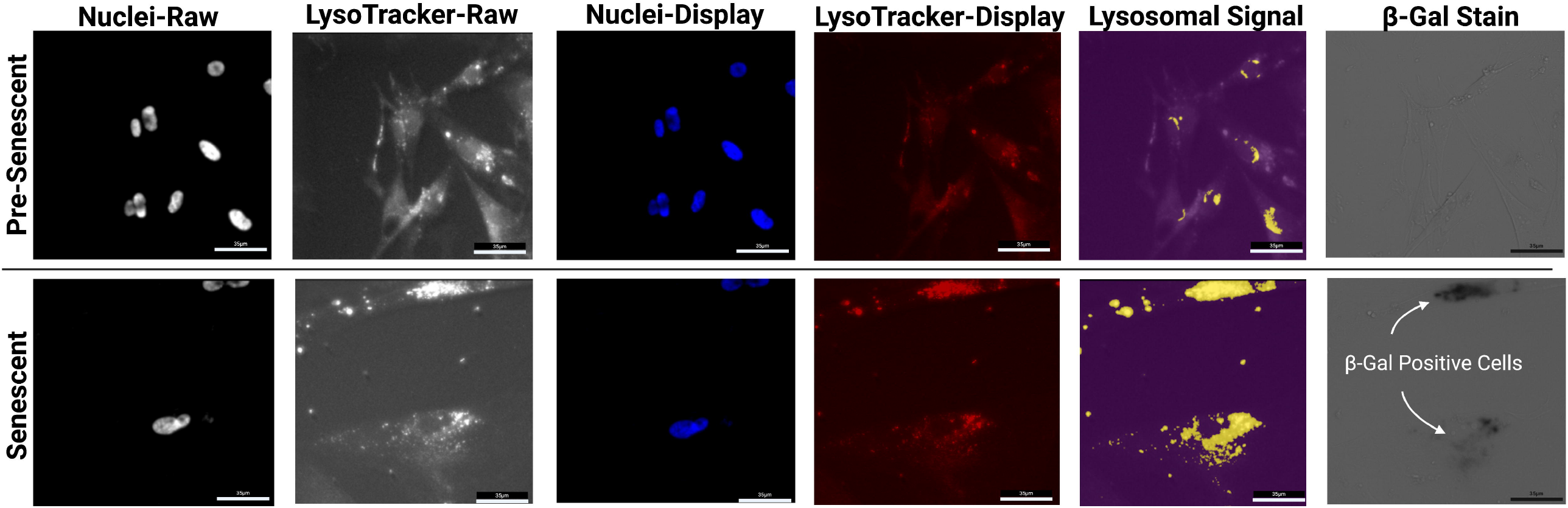
Representative live-cell fluorescence channels and post-fixation SA-β-Gal endpoint images in senescence-low and senescence-high IMR-90 cultures. Representative fields from pre-senescent (senescence-low) and senescent (senescence-high) IMR-90 cultures illustrate the appearance of the raw channels used for lysosomal profiling. Cells were stained live with LysoTracker Deep Red (lysosomal/acidic organelle signal) and Hoechst 33342 (live nuclear dye) and imaged by fluorescence microscopy using consistent acquisition settings within an experiment. Senescence-high cultures show brighter and more spatially expanded LysoTracker-enriched regions compared to senescence-low controls. Where shown, SA-β-Gal images are acquired after fixation and staining and are included as an orthogonal endpoint reference rather than a live-cell readout. Scale bars are shown in each panel; acquisition settings were kept constant within an experiment for comparability. Created in BioRender. Estrada, J. (2026) https://BioRender.com/w4zl4es

**Figure 3.**
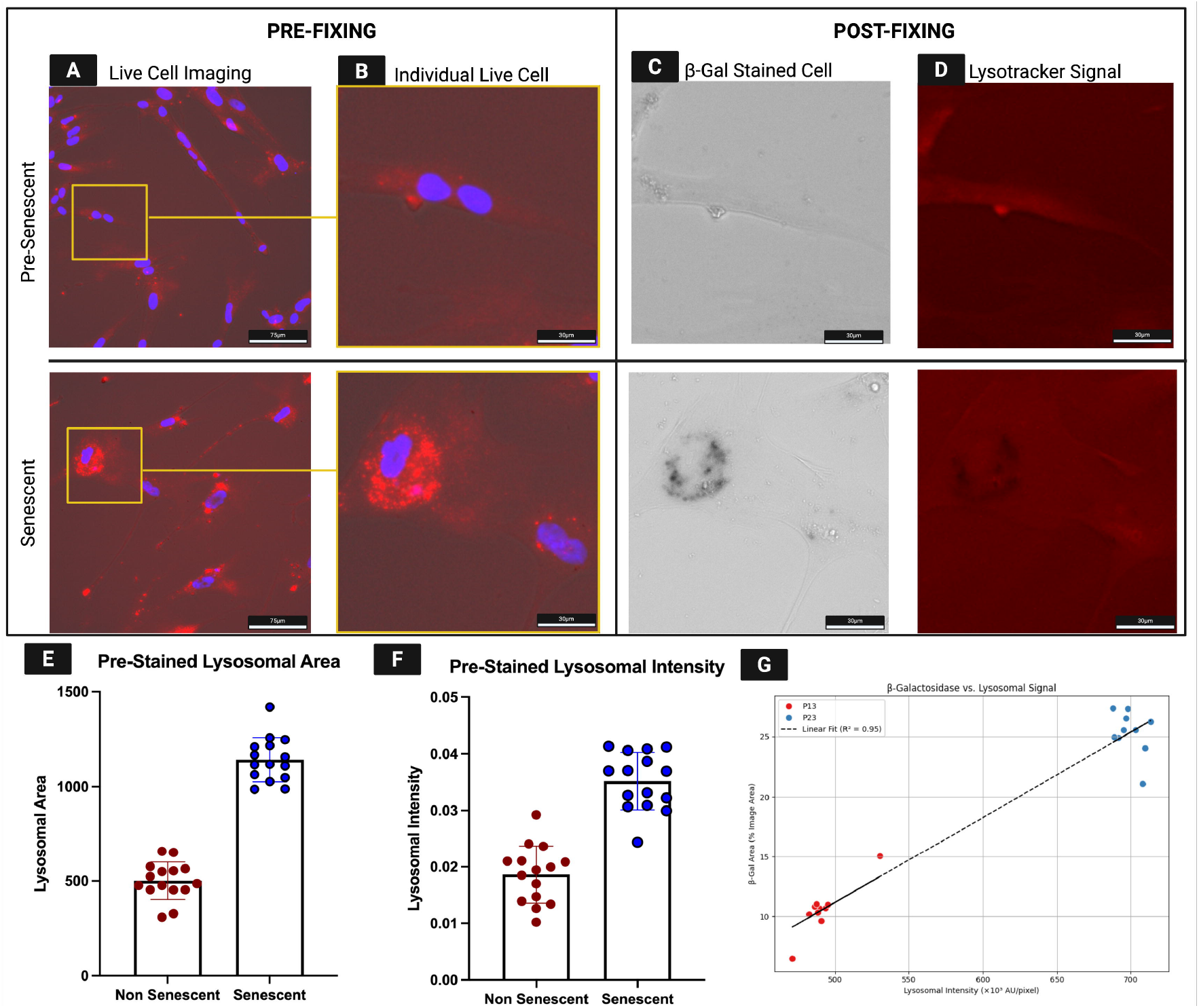
Linking live-cell LysoTracker signal to post-fixation SA-β-Gal staining across matched fields and replicate wells. Representative images show the same assay field before fixation (live staining) and after fixation and SA-β-Gal development, comparing senescence-low (pre-senescent) and senescence-high (senescent) IMR-90 cultures. **(A)** Live-cell overlay of the nuclear dye (blue) and LysoTracker Deep Red (red) prior to fixation. **(B)** Zoomed view of the boxed region in (A), highlighting a representative cell-level LysoTracker pattern under live imaging conditions. Following live imaging, cells were fixed and processed for SA-β-Gal. **(C)** Brightfield image after SA-β-Gal staining, illustrating the endpoint β-galactosidase signal in the matched field. **(D)** The LysoTracker channel from the corresponding field is shown for visual comparison with the SA-β-Gal endpoint readout. Quantification was performed using matched fields imaged before and after fixation. **(E)** Pre-fixation lysosome-enriched area per well, comparing senescence-low and senescence-high conditions. **(F)** Pre-fixation LysoTracker intensity per well, comparing the same conditions. In (E–F), each point represents one well (biological replicate), summarized from the stitched field(s) acquired for that well using the same analysis approach across conditions. **(G)** Correlation between pre-fixation LysoTracker readouts and post-fixation SA-β-Gal signal across wells. For each well, the SA-β-Gal-positive area (endpoint) was quantified from the post-fixation brightfield image and compared to the matched live LysoTracker measurement from the same field(s). Linear regression is shown with the corresponding goodness-of-fit value. Created in BioRender. Estrada, J. (2026) https://BioRender.com/8987vsq

4.3 Fix cells using the fixation solution provided in the SA-β-Gal staining kit prepared at the recommended 1x working concentration containing 2% formaldehyde and 0.2% glutaraldehyde. Incubate for 10 min to 15 min at room temperature.

4.4 Remove fixative and rinse wells twice with phosphate-buffered saline. Add SA-β-Gal staining solution at pH 6.0 containing X-gal substrate to each well.

4.5 Seal the plate to limit evaporation during incubation. Place the plate in a 37 °C dry incubator without CO_2_ for 12 h to 16 h.

4.6 Inspect wells by brightfield microscopy and document the staining pattern. Acquire representative images for each condition and quantify the fraction of SA-β-Gal-positive cells per well when reporting validation results.

CAUTION: Formaldehyde and glutaraldehyde are toxic and volatile. Prepare and use fixation solution in a chemical fume hood and wear appropriate personal protective equipment including gloves, eye protection, and a lab coat.

NOTE: If SA-β-Gal staining shows high background in the senescence-low control or weak staining in the senescence-high condition, repeat validation after adjusting plating density, fixation duration, incubation duration, and temperature consistency while keeping buffer pH at 6.0.

## REPRESENTATIVE RESULTS

Applying this protocol to IMR-90 fibroblasts across senescence-low and senescence-high conditions yields quantitative lysosomal readouts that can be summarized at the single-cell level and, when desired, at the whole-field level using stitched images. In the representative dataset shown here, stitched fluorescence fields were analyzed for each condition to provide a rapid summary of assay performance, including the number of nuclei detected from the nuclear channel and the fraction of the field occupied by LysoTracker-positive signal. These stitched-field summaries complement per-cell measurements by providing a direct, interpretable view of whether lysosomal remodeling differs between conditions under a fixed staining and imaging workflow.

A positive outcome is characterized by a visible and quantitative shift in lysosomal signal between senescence-low controls and senescence-high conditions. In the LysoTracker Deep Red channel, senescence-high cells typically show brighter and more spatially expanded perinuclear lysosome-enriched regions, whereas senescence-low controls show lower overall signal and more discrete punctate structures. In stitched-field summaries, this appears as condition-dependent changes in LysoTracker-positive area fraction and in mean intensity within the LysoTracker-positive mask, together with differences in per-nucleus normalized values when LysoTracker-positive area and intensity are expressed relative to the number of nuclei detected in the corresponding nuclear image (Table 1; Table S1).

**Table 1.** Whole-field lysosomal readouts from stitched images. Whole-field quantification from representative stitched fluorescence images stained with a live nuclear dye (nuclei detection) and LysoTracker Deep Red (acidic organelles). For each stitched field, nuclei were counted from the nuclear channel and LysoTracker-positive area was estimated by intensity thresholding in the LysoTracker channel. Mean intensity within the LysoTracker-positive mask and integrated intensity within the mask are reported in raw units. Per-nucleus values are calculated by dividing whole-field LysoTracker-positive area or integrated intensity by the nuclei count for the corresponding stitched field.

Stitched-field summaries are especially useful as a run-level check that staining quality, exposure settings, and plating density are within an interpretable range before scaling to larger collections of fields or downstream per-cell analysis. When SA-β-Gal staining is performed on parallel or matched wells, senescence-high conditions show a higher fraction of SA-β-Gal-positive cells than senescence-low controls. Agreement between elevated LysoTracker-based lysosomal remodeling and increased SA-β-Gal positivity provides a straightforward validation that the assay is capturing a senescence-associated phenotype. For experiments that include senescence induction strength or duration as variables, stronger or longer triggers yield higher lysosomal readouts and higher SA-β-Gal positivity, while milder conditions show intermediate responses.

Suboptimal experiments show characteristic signatures at the image and feature levels. Weak LysoTracker staining, overconfluent cultures, and focus drift reduce the clarity of lysosome-enriched regions and compromise cell separation, resulting in noisier measurements and reduced separation between conditions. In these cases, per-cell feature distributions become broad and overlapping, and well-level summaries show compressed dynamic range. Plate-to-plate changes in imaging settings can also shift intensities and reduce comparability if acquisition parameters are not held constant.

Overall, successful execution of the protocol yields: i) LysoTracker and nuclear images with clear single-cell resolution, ii) reproducible shifts in lysosomal remodeling features between senescence-low and senescence-high conditions, iii) optional SA-β-Gal validation showing increased positivity in senescence-high wells, and iv) well-level summaries that allow quantitative comparison of senescence induction and treatment effects.

## FIGURE AND TABLE LEGENDS

**Table S1. Additional stitched-field quantification (supplementary)**

Additional representative stitched fields quantified using the same workflow described for Table 1.

## DISCUSSION

This protocol describes a live-cell imaging workflow that reads out senescence from lysosomal remodeling and uses that signal to optionally train and apply a simple classifier. Several steps are particularly critical for obtaining a reliable assay. First, the biological model itself must be well controlled. Induction of replicative senescence in IMR-90 fibroblasts requires careful tracking of population doublings and parallel maintenance of early passage cultures as the non-senescent reference.^2,3,19^ Small deviations in passage history or confluence can shift the fraction of senescent cells and therefore change the dynamic range of the assay. Second, lysosomal and nuclear staining conditions must be tightly standardized. LysoTracker concentration, incubation time, and temperature, together with nuclear stain exposure, all affect signal intensity and segmentation quality. Finally, imaging settings and plate handling should be kept stable across runs so that differences in fluorescence reflect biology rather than focus drift or inconsistent acquisition. When these steps are consistent, lysosomal measurements separate senescence-low and senescence-high conditions in a way that aligns with conventional senescence markers such as SA-β-Gal, which reflects increased lysosomal mass during replicative aging.^10–13^

The protocol includes an optional branch that links live-cell lysosomal remodeling to SA-β-Gal validation. This branch introduces additional experimental sensitivities that are primarily practical rather than computational. The key requirement is that the same fields are imaged before and after SA-β-Gal staining so that lysosomal phenotypes are compared against the corresponding validation signal. Any mismatch between live and post-fixation fields can mix measurements across different cells and weaken the agreement between lysosomal remodeling and SA-β-Gal. In practice, this is addressed by careful plate handling and consistent imaging of matched fields, with spot checks to confirm that representative fields remain comparable throughout the workflow. Ground truth quality is equally important. SA-β-Gal staining must be tuned so that senescent cells develop a clear blue signal without excessive background in senescence-low cultures, which is sensitive to incubation time and the pH 6 staining conditions.

^12,13^ For this reason, users should confirm that SA-β-Gal conditions reproduce the expected differences in their hands before scaling up data collection.^10,12,13^ In parallel, visual inspection of raw images and overlays remains an essential quality step, because senescent fibroblasts often spread and overlap, and errors in separating adjacent cells or capturing the relevant cytoplasmic region can bias lysosomal measurements.^16^

A major strength of the workflow is its flexibility for different experimental questions, including senescence induction and senolytic testing. Users are not restricted to replicative aging as the only source of senescence. The same structure can be applied to senescence induced by irradiation, genotoxic stress, or oxidative stress, provided that appropriate controls and a measurable senescent fraction are included.^2–4,17,18^ Replicative senescence can still serve as a baseline reference because it is a well-established, biologically grounded model that captures the progressive nature of senescence acquisition and its links to lysosomal expansion.^10-13,19^ For treatment studies, senescence-low controls can be paired with induced senescence conditions, and candidate senolytics or senescence-modulating compounds can be evaluated by shifts in lysosomal remodeling and the resulting senescence burden. Additional channels may also be integrated in a bench-forward way, for example a cytoplasmic stain to improve cell boundary definition or a mitochondrial probe to capture complementary organelle changes, consistent with established approaches for assessing lysosome morphology and function.^6,16^ From a measurement perspective, the protocol is modular: users can begin with a minimal set of interpretable lysosomal features (area, intensity, and puncta-related measurements when resolvable) and expand only if needed.^16,20^

Several limitations should be acknowledged. The method centers on lysosomal remodeling and therefore assumes that increased lysosomal signal and lysosome-rich cellular morphology track with senescence. This is consistent with classic observations that SA-β-Gal reflects increased lysosomal mass during replicative aging and with newer work highlighting lysosome-centered mechanisms in senescence.^6–13^ However, lysosomal content varies across cell types and contexts, and lysosome-rich states may occur in cell types with high baseline lysosomal activity. In such settings, lysosomal remodeling may be less specific and may need to be interpreted alongside additional senescence markers. The validation labels themselves rely on SA-β-Gal, which is widely accepted but not infallible and can produce false positives under some conditions.^5,12,13^ In addition, like most imaging-based assays, the quantitative readout depends on consistent staining and acquisition conditions. When microscopes, magnifications, or exposure settings differ substantially, users should treat results as relative within a run and consider including local controls and validation rather than assuming direct transferability across platforms. Finally, the current implementation focuses on two-dimensional monolayers and does not capture the full complexity of tissue context or three-dimensional microenvironments.^12^

Despite these constraints, the protocol fills a practical gap between conventional senescence assays and more complex imaging workflows. Traditional readouts such as SA-β-Gal histochemistry, p16 or p21 immunostaining, EdU incorporation, and SASP measurements provide valuable information, but they are often endpoint, labor-intensive, and challenging to standardize across experiments.^3–5,13^ Recent work has shown that senescence can be recognized from nuclear morphology and other imaging features using machine learning, supporting the idea that image-derived measurements can provide robust senescence biomarkers.^21–23^ In this protocol, the emphasis remains on accessible vital dyes and direct, interpretable lysosomal measurements anchored to an established validation marker.^10–13^ Once an assay baseline is established and validated under consistent conditions, new plates require only a short LysoTracker plus nuclear staining step followed by routine imaging, after which senescence burden can be summarized at the cell or well level. This makes the workflow suitable for monitoring culture health over time, comparing senescence load under different oxygen tensions or stress conditions, and evaluating candidate senolytics or senescence-modulating compounds in modest-scale screening formats. In this way, the protocol provides an experimentally accessible route to more objective and reproducible senescence quantification while staying grounded in standard cell culture and imaging practices.

## ACKNOWLEDGMENTS

The research was funded by NIH grants R15CA274603 to JAM and TJB, R56DE033253 to ML, ST and JAM., and R01GM125870, 1R41AG081123 to ST and JAM.

